# Prompting Beyond Pairs: Decoupled Semantic Supervision for Knowledge-Guided Multiplex Virtual Staining

**DOI:** 10.64898/2026.07.31.741995

**Authors:** Yuqi Hu, Jiahao Wang, Kexiao Zheng, Yeying Jin, Hanry Yu

## Abstract

Virtual staining provides a non-invasive alternative to fluorescence microscopy, yet existing deep learning approaches fundamentally rely on pixel-aligned, multiplexed fluorescence targets for supervision. This dependence on rigidly paired data limits scalability, constrains flexibility in generating diverse subcellular structures, and becomes impractical in data-scarce biological settings. In this work, we introduce a semantic supervision paradigm for virtual staining, demonstrating that domain-knowledge prompts can effectively replace conventional pixel-level supervision. Unlike existing methods constrained by rigidly paired multiplex targets, our framework leverages biological prompts to decouple structural guidance from image translation. This decoupling enables high-fidelity, independent synthesis of multiple subcellular structures using only single-channel data. To ensure high-fidelity generation under weak supervision, we integrate self-supervised representation learning to mitigate data scarcity and incorporate direct preference optimization to suppress structural artifacts. Evaluations on the JUMP benchmark demonstrate that our approach effectively balances flexibility and fidelity, outperforming supervised baselines with a 43.3 % reduction in Average FID and an Average PCC of 0.912, while exhibiting high robustness in channel-deficient scenarios. Furthermore, the model generalizes across four in-house datasets to successfully multiplex six subcellular structures, overcoming the physical constraints of conventional fluorescent staining.

## 1 Introduction

Conventional fluorescent staining, though essential for subcellular profiling, suffers from phototoxicity and high costs that preclude longitudinal live-cell studies [7]. To overcome this, virtual staining provides a non-invasive computational alternative by predicting fluorescence directly from label-free brightfield images. Deep learning has significantly advanced this field; for instance, Cytoland [8] uses UNeXt2 architectures for generalizable marker prediction, while generative frameworks like DiffStain [3] employ mask-guided diffusion and spectral clustering to achieve multi-channel virtual staining with high structural fidelity.

While recent virtual staining methods show strong performance, they fundamentally rely on pixel-aligned, multiplexed fluorescence targets for supervision. Under this rigid supervision paradigm, comprehensively profiling diverse sub-cellular structures (e.g., actin, mitochondria, nuclei) necessitates exhaustively paired training data, where all target structures must be captured simultaneously in the ground truth. However, acquiring such strictly pixel-aligned, highly multiplexed data is often infeasible due to physical constraints like spectral overlap and phototoxicity. As a result, the scarcity of comprehensively multiplexed data severely limits the volume of usable training sets, ultimately bottlenecking model capacity and degrading generative performance. Furthermore, unavoidable domain shifts across diverse cell lines [14] render the acquisition of standardized datasets for every new experiment impractical. Therefore, there is a critical need for a flexible virtual staining paradigm capable of adapting to varying channel requirements without relying on rigidly paired, multiplexed supervision.

To address these limitations, we introduce a novel semantic supervision paradigm for multiplex virtual staining, reformulating the task as text-conditioned generation. Unlike existing methods, our framework leverages domain-knowledge through biological prompts to effectively replace conventional pixel-level supervision. This strategy decouples target structural specification from the image translation process. By doing so, we eliminate the fundamental reliance on rigidly paired, multiplexed pixel data and enable the flexible synthesis of diverse sub-cellular structures using independent channels. To ensure high structural fidelity under this weak supervision and resolve the inherent flexibility-fidelity trade-off, we integrate a morphology-aware self-supervised representation learning (SSL) module to mitigate data scarcity, alongside an adaptation of direct preference optimization (DPO) to explicitly enforce accurate morphological alignment and suppress artifacts. We rigorously evaluated this unified approach on the public JUMP Cell Painting benchmark[1] and multiple independent in-house datasets.

Our main contributions are as follows:

1. Semantic Supervision Paradigm: We propose a text-conditioned virtual staining framework that uses biological descriptive prompts instead of multiplexed pixel supervision, enabling the flexible, independent synthesis of diverse subcellular components from decoupled channel data.
2. High-Fidelity Synthesis Architecture: We integrate a morphology-aware SSL module and a biomedical-adapted DPO mechanism to explicitly enforce structural alignment under weak supervision. This unified approach overcomes the inherent limitations of decoupled training, achieving a 43.3 % reduction in Average FID compared to standard supervised baselines.
3. Robustness and Cross-Domain Generalization: We demonstrate the model’s remarkable resilience under channel-deficient scenarios (maintaining an Average SSIM of 0.923 and FID of 34.69) and validate its practical generalizability by successfully multiplexing six subcellular structures across four independent, inhouse cell-line datasets.

## 2 Methods

Let 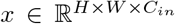 denote the input brightfield image. Conventional virtual staining models are restricted to a predetermined fixed output dimensionality, mapping *x* to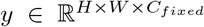. This rigidity necessitates training distinct models whenever the target organelle or channel count changes. To overcome this limitation, we reformulate virtual staining as a text-conditioned generation task. We introduce a textual prompt *t* to semantically specify the target component, enabling our model *M*_*θ*_ to learn the conditional distribution *p*(*y*|*x, ∈*(*t*)), where *∈* is a text encoder. Unlike hard-coded architectures, this design dynamically defines output semantics via the prompt, allowing a single unified model to generate diverse staining modalities (e.g., {*y*_*mito*_, *y*_*nuclei*_, …}) from the same input *x*. To effectively train *M*_*θ*_ in practical scenarios with limited and imbalanced training data across different organelle types, we propose a three-stage training strategy (Fig. 1). This strategy progressively integrates self-supervised learning (SSL) [6] for robust feature extraction and direct preference optimization (DPO) [17] for aligning generated outputs with desired staining quality, as detailed in Section 2.2 and 2.3.

**Fig. 1.**
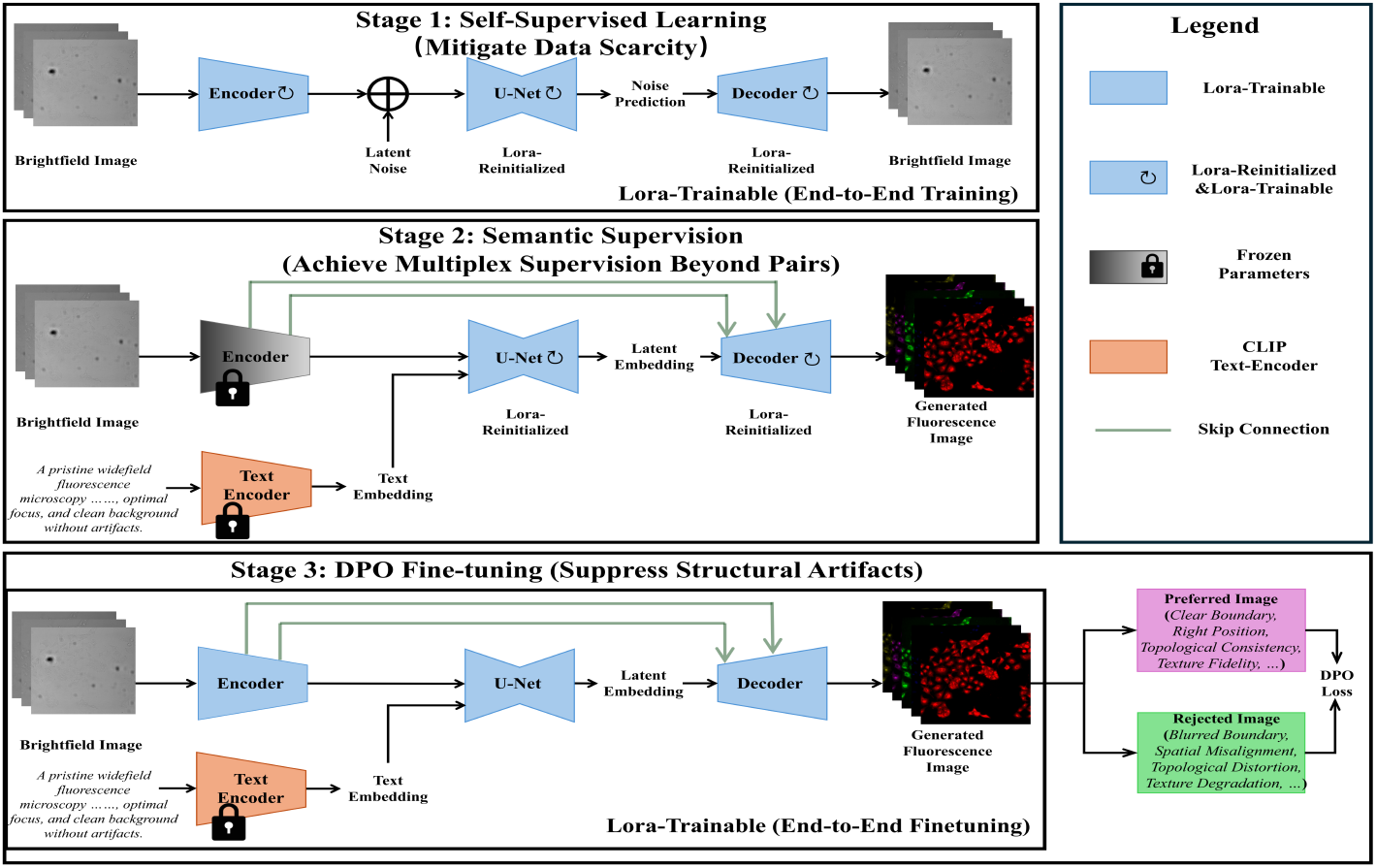
Proposed training strategy. Figure 1. Three-Stage Framework for Decoupled Virtual Staining. (Stage 1) Self-Supervised Pre-training: The encoder is pre-trained via brightfield in-painting (*x→ x*) without skip connections to mitigate data scarcity and learn robust semantic representations. (Stage 2) Prompt-Guided Decoupled Adaptation: Encoder weights are transferred, and biological domain-knowledge prompts are introduced to decouple structural guidance from image translation (*x→ y*). Skip connections are integrated to recover spatial details for high-fidelity synthesis. (Stage 3) DPO Fine-tuning: The entire model is aligned using Direct Preference Optimization (DPO) to suppress structural artifacts and ensure overall generation fidelity.

### 2.1 Text-Conditional Generative Architecture

Our virtual staining model is built on Stable Diffusion Turbo (SD-Turbo) and follows the pix2pix-Turbo [9] paradigm for conditional image-to-image translation (Fig. 1, Stage 2). Architecturally, we adopt the latent diffusion backbone [12] with an encoder–U-Net–decoder pipeline, and fine-tune it in a parameter-efficient manner using low-rank adaptation (LoRA) [5] modules. To ensure high-fidelity generation, we use encoder-decoder skip connections [13] to propagate high-frequency features. This preserves the structural integrity of fine-grained patterns, which is critical for the independent synthesis of multiple subcellular structures using only single-channel data.

#### Domain-Knowledge Integration via Descriptive Prompts

To replace conventional pixel-level supervision and control synthesis *p*(*y*|*x, ∈*(*t*)), we ground our model in wet-lab domain knowledge. Comprehensive experimental descriptors (e.g., cell line, treatment, optics) and representative fluorescent images are processed by Gemini [15] to distill visual phenotypes into standardized biological prompts (*t*_*std*_) for each subcellular structure. To prevent linguistic overfitting, each *t*_*std*_ is augmented into *K* = 5 semantic variants. The selected prompt *t* is embedded via CLIP [10] into *∈*(*t*) and injected into the U-Net via cross-attention, providing semantic supervision to guide the generation of the target modality *y*.

### 2.2 Self-Supervised Representation Learning

Since paired fluorescence data is scarce, we first exploit abundant unlabeled brightfield images for representation pre-training via a latent denoising autoen-coder (l-DAE) framework [2] (Stage 1). Crucially, during this pre-training phase, we deliberately remove the encoder–decoder skip connections described in Section 2.1. By eliminating direct spatial shortcuts, we impose an information bottleneck [16] that precludes trivial identity mapping and compels the encoder to learn compact, high-level semantic representations of biological structures to solve the reconstruction task (*x → x*).

### 2.3 Preference Alignment via DPO Fine-tuning

To enhance generative fidelity for complex organelles, we apply Direct Preference Optimization (DPO) [11, 17]. Using training triplets (*x, y*_*w*_, *y*_*l*_)—where *x* is the input, *y*_*w*_ is the high-fidelity ground truth fluorescence (chosen target), and *y*_*l*_ is a manually curated suboptimal output (rejected target)—we define an implicit reward based on the negative Mean Squared Error (MSE). Specifically, the reward margin for the active policy model *θ* is formulated as the difference in reconstruction errors:

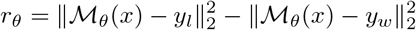

To stabilize learning and preserve structural integrity, we optimize against a frozen reference model *θ*_ref_ and regularize the DPO objective with pixel-wise (*L*_MSE_) and perceptual (*L*_LPIPS_) losses [19]:

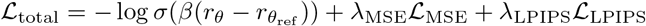

where *σ* is the sigmoid function and *β* controls the deviation from the reference model.

## 3 Experiments & Results

### 3.1 Experimental Details

#### Datasets

We evaluated our framework by training and testing it on two datasets. First, we used the public JUMP Cell Painting dataset (cpg0000) [1]^4^. Following DiffStain [3], we removed out-of-focus samples and split the data by plate (nine for training, one for testing), yielding 45,000 training and 9,000 testing paired images per channel (DNA, ER, RNA, AGP, Mito). Second, to demonstrate the framework’s adaptability in diverse biomedical research scenarios, we compiled an in-house dataset comprising four cell lines (3T3, HepaRG, HFF, HUVEC). To match the JUMP protocol’s three-channel brightfield input, we concatenated the optimal focal plane with the top and bottom z-layers. Corresponding ground truth targets (actin, nucleus, mitochondria, ER, RNA, and AGP) were derived using Maximum Intensity Projections (MIP) from z-stacks.

#### Implementation Details

Our framework was implemented in PyTorch and trained on two NVIDIA RTX 4090 GPUs. The self-supervised pre-training stage was initialized using SD-Turbo weights^5^.

#### Evaluation Metrics

Staining fidelity was quantified using Mean Absolute Error (MAE) and Pearson Correlation Coefficient (PCC) for pixel-level accuracy, Structural Similarity Index Measure (SSIM) [18] for structural preservation, and Fréchet Inception Distance (FID) [4] for perceptual domain alignment. During the evaluation phase, we strictly applied a leave-one-out evaluation strategy on the test set to compute all metrics.

### 3.2 Results

To validate the effectiveness of our proposed framework, we conducted a comprehensive ablation study using a vanilla Stable Diffusion model with null prompts as the baseline. We also benchmarked our method against DiffStain, a competitive model trained on the same public dataset. As shown in Table 1 and Fig. 2, we incrementally integrated our proposed modules to assess their individual contributions. We first introduced Simple Prompts (SP), utilizing basic class-level keywords (e.g., ‘mitochondria’ or ‘nucleus’) to distinguish between subcellular components. This was followed by the integration of biological descriptive prompts, Self-Supervised Learning (SSL), and Direct Preference Optimization (DPO). Both qualitative and quantitative results demonstrate that each component progressively enhances average image quality. Notably, our final method surpasses DiffStain across all available metrics, confirming the efficacy of the proposed modules.

**Table 1.** Ablation Study on prompts and training strategies. **SP**: Simple Prompts. **DP**: Descriptive Prompts. **SSL**: Self-supervised Learning. **DPO**: Direct Preference Optimization. **ER**: Endoplasmic Reticulum. **Mito**: Mitochondria.

| Method | Average |  |  |  | RNA |  |  |  | ER |  |  |  |
| --- | --- | --- | --- | --- | --- | --- | --- | --- | --- | --- | --- | --- |
|  | MAE↓ | PCC↑ | SSIM↑ | FID↓ | MAE↓ | PCC↑ | SSIM↑ | FID↓ | MAE↓ | PCC↑ | SSIM↑ | FID↓ |
| DiffStain | 0.088 | 0.862 | 0.636 | – | 0.076 | 0.914 | 0.696 | – | 0.093 | 0.881 | 0.655 | – |
| Baseline | 0.028 | 0.818 | 0.885 | 51.68 | 0.028 | 0.870 | 0.888 | 30.96 | 0.027 | 0.818 | 0.882 | 35.29 |
| + SP | 0.024 | 0.875 | 0.913 | 42.21 | 0.024 | 0.925 | 0.921 | 26.60 | 0.023 | 0.897 | 0.918 | 32.33 |
| + DP | 0.024 | 0.879 | 0.914 | 34.11 | 0.024 | 0.930 | 0.920 | 22.84 | 0.022 | 0.900 | 0.918 | 26.30 |
| + DP + SSL | 0.020 | 0.906 | 0.929 | 30.30 | 0.022 | 0.946 | 0.934 | 24.37 | 0.020 | 0.927 | 0.936 | 24.20 |
| + DP + SSL + DPO | **0.019** | **0.912** | **0.933** | **29.31** | **0.017** | **0.951** | **0.944** | **20.89** | **0.016** | **0.932** | **0.943** | **22.44** |

| Method | Mito |  |  |  | AGP |  |  |  | Nucleus |  |  |  |
| --- | --- | --- | --- | --- | --- | --- | --- | --- | --- | --- | --- | --- |
|  | MAE↓ | PCC↑ | SSIM↑ | FID↓ | MAE↓ | PCC↑ | SSIM↑ | FID↓ | MAE↓ | PCC↑ | SSIM↑ | FID↓ |
| DiffStain | – | – | – | – | – | – | – | – | – | – | – | – |
| Baseline | 0.040 | 0.797 | 0.854 | 76.33 | 0.030 | 0.817 | 0.867 | 41.29 | 0.016 | 0.786 | 0.934 | 74.53 |
| + SP | 0.036 | 0.856 | 0.886 | 82.34 | 0.022 | 0.874 | 0.898 | 39.80 | 0.015 | 0.822 | 0.942 | 30.01 |
| + DP | 0.035 | 0.876 | 0.893 | 55.88 | 0.023 | 0.872 | 0.897 | 37.02 | 0.015 | 0.823 | 0.941 | 28.52 |
| + DP + SSL | **0.028** | 0.902 | **0.913** | 53.87 | **0.019** | 0.899 | 0.909 | **24.81** | 0.020 | 0.854 | 0.954 | 24.23 |
| + DP + SSL + DPO | 0.030 | **0.908** | 0.910 | **53.86** | 0.020 | **0.901** | **0.911** | 27.10 | **0.010** | **0.869** | **0.959** | **22.27** |

**Fig. 2.**
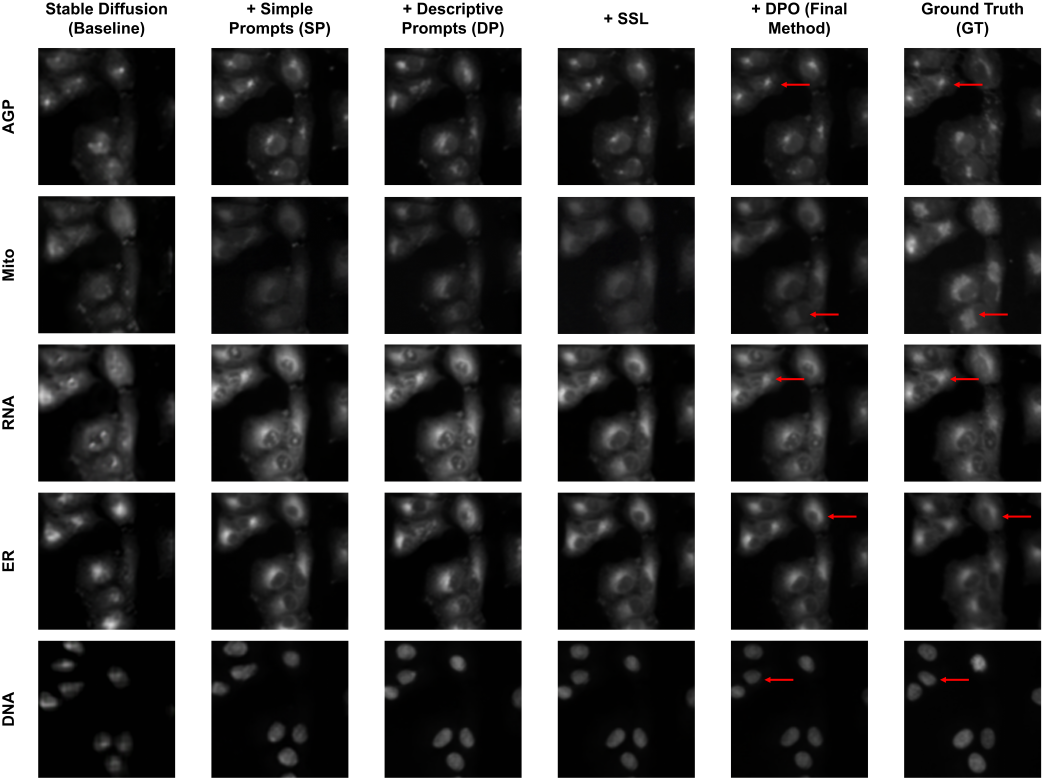
Visual comparison of ablation study results. Columns show the progressive improvement of generated images. “Stable Diffusion” corresponds to the Baseline in Table 1. Subsequent columns show the addition of Simple Prompts (SP), Descriptive Prompts (DP), Self-supervised Learning (SSL), and Direct Preference Optimization (DPO), respectively. The final column is the Ground Truth (GT). Red arrows high-light details improved by the proposed components.

To evaluate the framework’s applicability in real-world biological workflows where full-channel acquisition is often inconsistent, we simulated data scarcity scenarios by training on channel-deficient subsets. We designed two mixed-modality training settings: (1) a single-channel depletion setting (A+B), where the model was trained on a combined dataset comprising Group A (lacking RNA) and Group B (lacking ER); and (2) a severe sparsity setting (C+D+E), where the training set consisted of three groups, each lacking two specific modalities (e.g., Group C lacking RNA and ER, Group D lacking RNA and mitochondria, and Group E lacking ER and mitochondria). All models were evaluated on a fully-paired test set. As presented in Table 2, our method demonstrates remarkable robustness under data scarcity: models trained on incomplete data (A+B and C+D+E) exhibit only marginal performance degradation and consistently out-perform both the Baseline and DiffStain methods. This confirms our strategy’s efficacy in fully exploiting partial data, proving its high viability for wet-lab scenarios characterized by inconsistent channel acquisition.

**Table 2.**
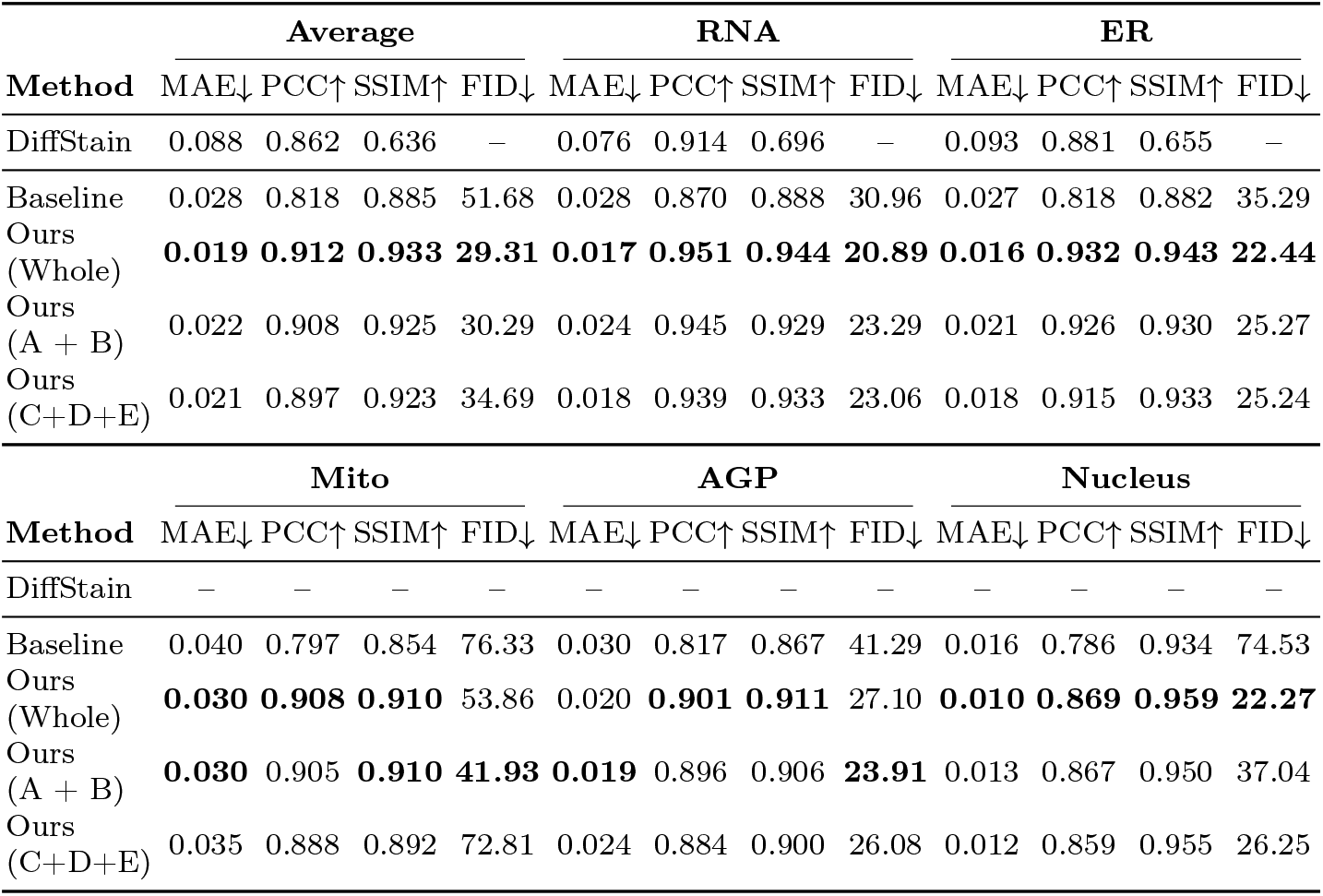
Quantitative comparison of dataset composition strategies. **Whole**: Complete dataset training. **A+B**: Training with subsets missing single channels (A: w/o RNA, B: w/o ER). **C+D+E**: Training with subsets missing two components each. Values indicate our method maintains high performance even with incomplete data.

Furthermore, as illustrated in Fig. 3, we demonstrated the adaptability of our proposed framework by training it on four distinct in-house datasets, corresponding to diverse cell lines (3T3, HepaRG, HFF, and HUVEC). The proposed method simultaneously multiplexes six distinct subcellular components, thereby overcoming the inherent constraints of conventional staining protocols and optical microscopy. The framework demonstrates robust virtual staining performance across significant variations in cellular morphology, culture density, and subcellular organization, confirming its practical viability for deployment in real-world biomedical research workflows.

**Fig. 3.**
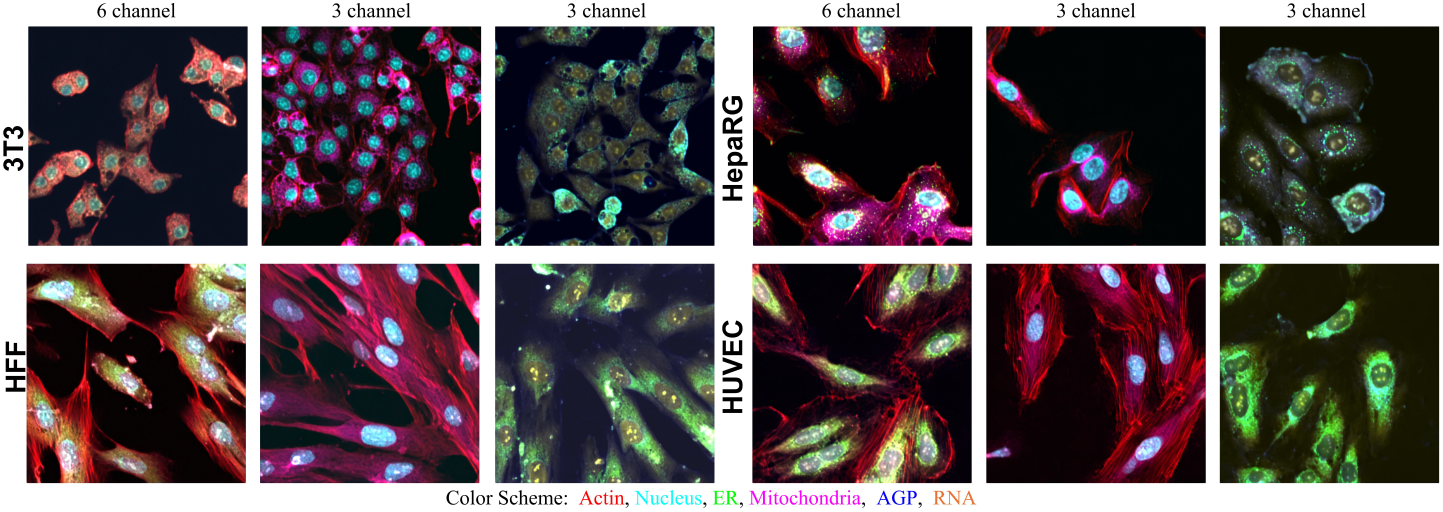
Multiplexed virtual staining via decoupled component generation. By effectively decoupling multiplexed components, our framework generates six simultaneous subcellular structures across four distinct datasets (3T3, HepaRG, HFF, and HUVEC) (first image of each group), outperforming conventional fluorescence microscopy which is limited to fewer components (second and third images).

## 4 Conclusion

We introduce a semantic supervision paradigm for virtual staining that eliminates the fundamental reliance on rigidly paired, multiplexed fluorescence tar-gets. By utilizing domain-knowledge prompts to separate semantic control from the image translation process, our framework enables the independent, high-fidelity synthesis of diverse subcellular structures using only single-channel data.

Furthermore, integrating self-supervised pretraining and DPO mitigates data scarcity and suppresses structural artifacts. By breaking the fixed-channel bottleneck and demonstrating robust generalization in channel-deficient scenarios, our prompt-guided approach establishes a highly scalable and adaptable reagent-free paradigm for subcellular profiling.

## Footnotes

4 https://open.quiltdata.com/b/cellpainting-gallery/tree/cpg0000-jump-pilot/source_4/images

5 https://huggingface.co/stabilityai/sd-turbo

